# A high-throughput screen identifies small molecule modulators of alternative splicing by targeting RNA G-quadruplexes

**DOI:** 10.1101/434647

**Authors:** Jing Zhang, Samuel E. Harvey, Chonghui Cheng

## Abstract

RNA secondary structures have been increasingly recognized to play an important regulatory role in post-transcriptional gene regulation. We recently showed that RNA G-quadruplexes, which serve as cis-elements to recruit splicing factors, play a critical role in regulating alternative splicing during the epithelial-mesenchymal transition. In this study, we performed a high-throughput screen using a dual-color splicing reporter to identify chemical compounds capable of regulating G-quadruplex-dependent alternative splicing. We identify emetine and its analog cephaeline as small molecules that disrupt RNA G-quadruplexes, resulting in inhibition of G-quadruplex-dependent alternative splicing. Transcriptome analysis reveals that emetine globally regulates alternative splicing, including splicing of variable exons that contain splice site-proximal G-quadruplexes. These data suggest the use of emetine and cephaeline for investigating mechanisms of G-quadruplex-associated alternative splicing.

## INTRODUCTION

RNA secondary structures have been shown to play key roles in gene expression through regulating alternative splicing [1–4], translational regulation [5, 6], and transcriptional termination [7]. One type of RNA secondary structures is RNA G-quadruplexes, which are formed within guanine-rich sequences. These structures can be formed within a single strand or between multiple strands of RNA, where four G-tracts of two or more guanines separated by short stretches of other nucleotides are assembled in layered loops bound together through Hoogsteen hydrogen bonding. Accumulating evidence has indicated that G-quadruplex structures are associated with many human diseases, including neurological disorders [8] and cancer [9, 10]. Although much effort has been directed toward understanding the importance of RNA G-quadruplexes in biology, few small molecules capable of targeting these structures to modulate biological functions have been discovered or developed.

Alternative splicing of pre-messenger RNA occurs in up to 95% human genes during development or in response to extracellular stimuli [11, 12]. During these processes, a single gene can produce protein isoforms with different, even opposing functions. Disruption of splice site recognition and misregulation of alternative splicing causes many human diseases, including autoimmune disorders [13], neurodegenerative diseases [14] and cancer progression [15, 16]. Therefore, alternative splicing is an important and potential target for the treatment of various types of human disorders. Alternative splicing is principally regulated by recruitment of RNA binding proteins to conserved cis-regulatory elements on the pre-mRNA. Some progress has been made in the discovery of small-molecules designed to selectively target splicing factors and alternative splicing events [17–19]. The identification of small-molecules targeting RNA secondary structures that are associated with disease-relevant alternative splicing, such as G-quadruplexes, has been very limited.

In this study, we utilized a dual-color splicing reporter containing a G-quadruplex secondary structure proximal to the splice sites that allowed us to measure changes in cassette exon inclusion and skipping dependent on the adjacent G-quadruplex. Using this splicing reporter, we performed a high-throughput screen to identify small molecules that affect G-quadruplex-dependent alternative splicing. We found that the analogous small molecules emetine and cephaeline regulate alternative splicing by disrupting G-quadruplex structures. We further demonstrate that these small molecules modulate splicing when G-quadruplexes are located either upstream or downstream of the cassette exon. In addition, we show that emetine-regulated alternative splicing affects G-quadruplex associated alternative splicing across the transcriptome. Therefore, our study identifies new small molecule compounds with the potential to regulate alternative splicing by targeting G-quadruplexes.

## MATERIALS AND METHODS

### Plasmids and primers

The bichromatic fluorescent reporters, CRQ, CRQ G4m2, CRQ G4m and scrambled sequences, were cloned into EcoRI and XhoI sites of the RG6 vector, which are located downstream of the variable exon [3]. To determine whether the location of the G-quadruplex affects alternative splicing, these sequences were also inserted into BsiW1 and Not1 sites of the RG6 vector, which are located upstream of the variable exon.

### Cell lines and generation of stable cell lines

To generate the stable cell lines expressing the bichromatic fluorescent reporters, CRQ and CRQ G4m reporters were transfected into HEK 293A cells, termed HEK 293A-CRQ and HEK 293A-CRQ G4m. Cells stably expressing the reporters were selected using G418 and inspected visually under a fluorescent microscope for EGFP and dsRed. After two weeks of selection, HEK 293A-CRQ were sorted for EGFP-positive cells and HEK 293A-CRQ G4m were sorted for dsRed-positive cells by flow cytometry for downstream screening.

### Cell culture

HEK 293FT, HEK 293A-CRQ and HEK 293A-CRQ G4m cells were maintained in Dulbecco’s Modified Eagle’s Medium (DMEM) supplemented with 10% fetal bovine serum (FBS), 100 units/ml penicillin and streptomycin, 2mM L-Glutamine in 5% CO_2_ at 37°C. Maintenance of HMLE-Twist and MCF10A were described previously [20, 21].

### High-throughput screening

The screen was performed on the ImageXpress Micro Confocal High-Content Imaging System. HEK 293A-CRQ cells (3,000 per well) were seeded into 384-well plates overnight prior to treatment with the MicroSource Custom Collection library. This library contains 2,700 compounds: 50% of the compounds are approved drugs, 35% are natural products and 15% are compounds with established and diverse bioactivity. HEK 293A-CRQ G4m cells, which solely express dsRed, were used as a positive control for dsRed in images. HEK 293A-CRQ cells that were treated with DMSO were used as a positive control for EGFP in images. Images of each well were recorded under a 10× microscope immediately after addition of the compounds (10 μM) as a day 0 timepoint. After 48 hrs of treatment, images were taken and analyzed to obtain the numbers of dsRed cells in each well. The library was screened in duplicate. Percentage of dsRed cells were calculated by the number of red colored cells in the fluorescence images divided by the total number of cells in the corresponding images. A cut-off with total cell number ≥ 600 and percent of red cells ≥ 7% was set to select for positive hits. After the screening, a dose response assay and an RT-PCR analysis were performed for each positive compound to filter out false positives.

### Reporter transfection assay

For reporter transfection assays, HEK 293FT (1×10^5^) cells were seeded in a 24-well plate prior to transfection. The bichromatic fluorescent reporters (800 ng) were transfected using Lipofectamine 2000 (Life Technologies). 24 hrs after transfection, small molecule compounds or DMSO were added into each well. Fluorescence images were taken under 10x microscopy after 48 hrs of drug treatment. RNA was extracted after 48 hrs of drug treatment to perform RT-PCR analysis of splicing.

### Circular dichroism spectroscopy

To conduct the circular dichroism spectroscopy measurement, RNA oligonucleotides, CRQ (5’-GGGCGGCGGGCGGGCUGGGG-3’) and I-8 (5’-GCUUUGGUGGUGGAAUGGUGCUAUGUGG-3’), were purchased from Integrated DNA Technologies and dissolved in Nuclease-free water at a concentration of 100 μM. RNA oligonucleotides (5 μM) were prepared in annealing buffer (10 mM Tris-HCl, pH=7.4 and 100 mM K^+^) and annealed by heating to 95°C and then cooling down slowly to room temperature. After annealing, different amount of emetine were added and the circular dichroism of RNA oligonucleotides was measured at 20°C by a Jasco J-815 spectropolarimeter. An average of five CD spectra ranging from 220 to 320 nm was collected at a bandwidth of 1 nm in a 0.1-mm path length cuvette, with a response time of 2 sec/nm and continuous scan mode.

### RNA extraction, semi-quantitative and real-time RT-PCR

Total RNA was isolated from cultured cells with total RNA kit (Omega Bio-tek). Isolated total RNA (125 ng) was used for reverse transcription with the GoScript Reverse Transfection System to produce cDNA (Promega). For semi-quantitative RT-PCR, Hot StarTaq DNA polymerase (QIAGEN) was used for no more than 30 cycles. The PCR products were separated on 2% agarose gels. The relative signal intensity of each band was obtained. For real-time PCR, Bio-Rad iQ SYBR Green Supermix was used on a BioRad CFX instrument.

### RNA sequencing and G-quadruplex prediction

RNA was extracted using Trizol from HMLE-Twist and MCF10A cells that were treated with DMSO or emetine (10 μM) for 24 hrs. Two biological replicates were performed per experimental condition. Poly-A selected RNA-seq libraries were generated using TruSeq Stranded mRNA Library Prep Kits (Illumina) and subjected to 100 bp PE stranded RNA sequencing on an Illumina Hiseq 4000. RNA-seq reads were aligned to the human genome (GRCh37, primary assembly) and transcriptome (Gencode v24 backmap 37 comprehensive gene annotation) using STAR v2.5.3a [22] with the following parameters: STAR --runThreadN 16 --alignEndsType EndToEnd --outSAMtype BAM SortedByCoordinate. Differential alternative splicing was quantified using rMATS v3.2.5 [23] with the following non-default parameters: -t paired -len 100-analysis U - libType fr-firststrand. Significant differentially spliced cassette exons were quantified using only unique junction reads and the following cutoffs: FDR < 0.05, delta PSI ≥ 0.1, average junction reads per cassette event per replicate ≥ 20. Predicted G-quadruplexes were curated from a strand-specific list of all sequences in the GRCh37 version of the human genome matching the regular expression GxNy, where N equals any series of nucleotides other than G, x ≥ 2 or 3 and 1 ≤ y ≤ 7 to identify all sequences with G-tracts of 2-3 or more separated by 1-7 of any other base. Bedtools v2.26.0 and custom bash and python scripts were used to intersect emetine-dependent cassette exons, the flanking variable exons, and the 250 nucleotides flanking each splice site with these G-quadruplexes. RNA-sequencing data has been deposited under accession number GSE113505.

## RESULTS

### The small molecule emetine inhibits G-quadruplex-dependent alternative splicing

We previously constructed a bichromatic fluorescent splicing reporter, which contains a cassette exon and a known G-quadruplex containing sequence, CRQ, inserted just downstream of the 5’ splice site [3]. The splicing reporter expresses EGFP as a readout of alternative exon inclusion and dsRed when exon skipping occurs (Figure 1A). CRQ is a 20-nucleotide (nt) RNA sequence that forms a bona-fide G-quadruplex secondary structure (Supplementary Figure S1A, and Ref. [3]). Cells expressing this reporter primarily produce the exon inclusion isoform and show EGFP fluorescence. Disrupting the G-quadruplexes in CRQ results in the production of the exon skipping isoform and shifts the fluorescence from EGFP to dsRed (Supplementary Figure S1B and Ref. [3]). To identify compounds that regulate alternative splicing by disrupting G-quadruplexes, we developed an imaging assay to quantify changes in the ratio of EGFP and dsRed signals in an HEK 293A cell line stably expressing the CRQ-splicing reporter. The predominant green fluorescence and minute level of red fluorescence in these cells provided an optimal system to identify compounds that shifts the splicing from the green inclusion isoform to the dsRed skipping isoform. We employed the MicroSource Custom Collection library that contained 2700 small molecule compounds and treated the CRQ-reporter expressing HEK 293A cells in 384-well plates. After 48 hrs of treatment, cells in individual wells were imaged by ImageXpress microscopy and monitored for changes in total cell number as well as an increase in the dsRed signal, which represents exon skipping events. After initial identification of 38 hits that showed increased percentage of red-colored cells (Supplementary Table S1), we conducted a secondary screen by dose-response assay and found 6 out of the 38 compounds that exhibited a dose-dependent increase in red signals (Supplementary Table S2). Moreover, 3 out of these 6 compounds (emetine, obtusaquinone, and Sanguinarine sulfate) showed effects on shifting splicing by RT-PCR analysis. We chose to study emetine, a natural alkaloid produced from ipecas root [24], because of its highest potency on inhibiting CRQ-dependent splicing (Figure 1B, 1C, Supplementary Figure S1C). Quantitative real-time PCR (qPCR) also confirmed that 20 μM emetine treatment caused a shift of alternative splicing to the skipped isoform (Figure 1C).

**Figure 1.**
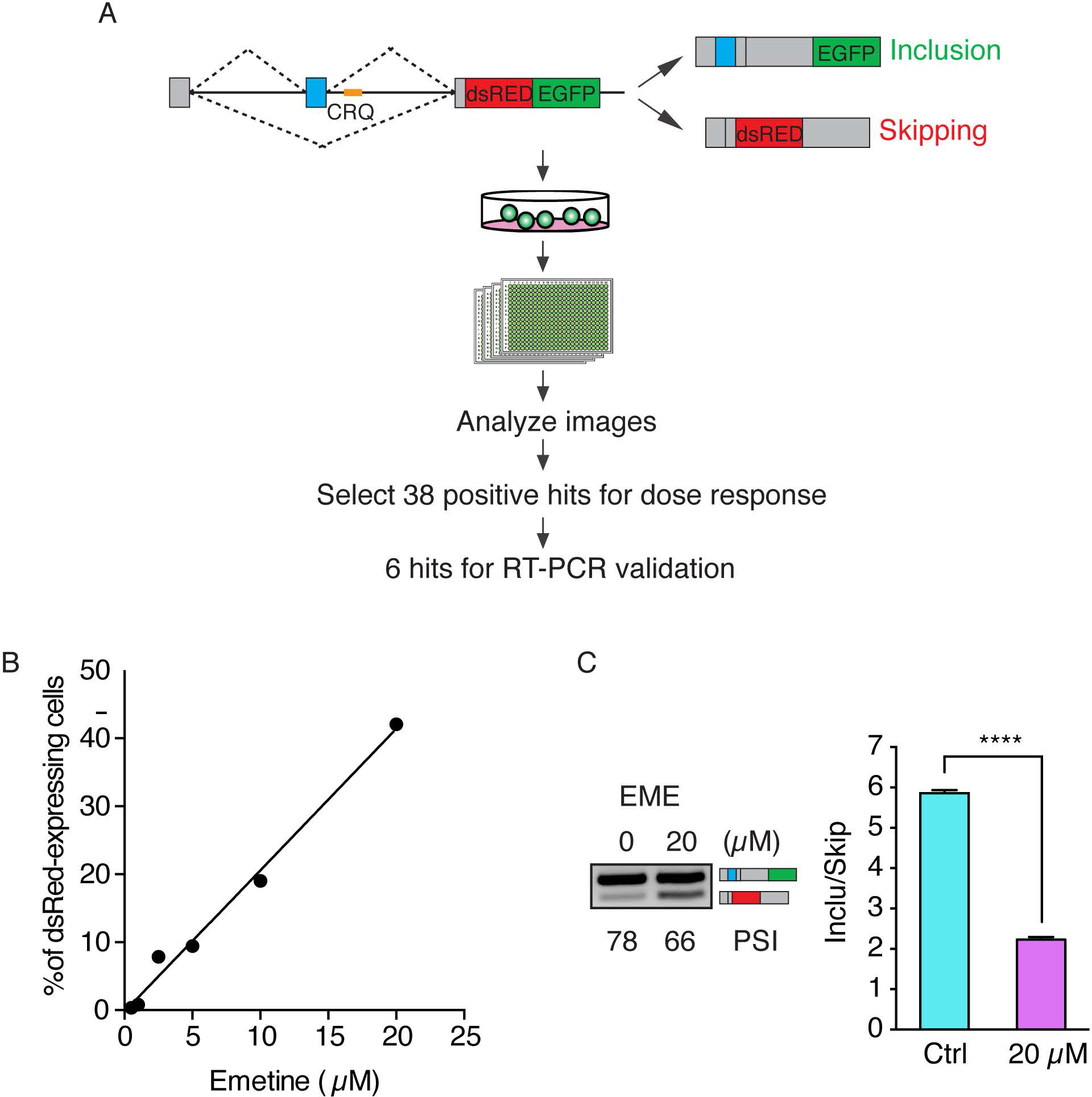
A high-throughput screen to identify modulators of G-quadruplex-dependent alternative splicing. (A) Flowchart of the high-throughput screening assay. The mini-gene splicing reporter is depicted on the top panel. The CRQ G-quadruplex sequence is colored in yellow. Inclusion of the variable exon (in blue) results in the production of EGFP. Skipping of the variable exon generates dsRed. (B) Dose response relationship between emetine and the percentage of dsRed-expressing cells. (C) Semi-quantitative RT-PCR (Left) and qRT-PCR analysis (Right) of emetine (20 μM) treatment on HEK 293A-CRQ for 48 hrs (Error bars represent S.E.M. ****, p < 0.0001; Student’s *t*-test).

### Emetine regulates alternative splicing by disrupting G-quadruplex secondary structure

To assess whether emetine inhibits exon inclusion by interfering with G-quadruplexes, we compared the effect of emetine in G-quadruplex containing CRQ minigene to the control non-CRQ minigene. As seen in Figure 2A-2C, control non-G-quadruplex containing minigene did not show any changes in splicing in response to emetine treatment. The G-quadruplex containing CRQ splicing minigene, which showed increased exon inclusion compared to the control minigene, showed a decrease in exon inclusion after emetine treatment, suggesting that the G-quadruplex-dependent increase in exon inclusion is inhibited by emetine. Furthermore, we introduced mutations disrupting the guanine tetrads in CRQ constructs to abolish the G-quadruplex structure (Supplementary Figure S1A). We disrupted the G-quadruplex structure in CRQ by substituting the middle two guanine nucleotides with two adenosines, which prevented the sequence from forming a G-quadruplex without dramatically altering the guanine composition of the sequence (Supplementary Figure S1A). This G-quadruplex mutant, termed CRQ G4m2, partially abrogated CRQ-mediated exon inclusion, resulting in a decrease in the ratio of exon inclusion to skipping (Figure 2A-2C-2C). Emetine treatment with this minigene showed minimal changes in alternative splicing, suggesting that emetine loses its inhibitory effect in exon inclusion in the absence of an intact G-quadruplex (Figure 2A-2C). Furthermore, disrupting the G-quadruplex by substituting all four GGG repeats (CRQ G4m), which completely deminished G-quadruplex-mediated exon inclusion (Figure 2A-2C), also abolished the effect of emetine-mediated inhibition on exon inclusion (Figure 2A-2C), once again indicating that the G-quadruplex is required for emetine-mediated inhibition of exon inclusion.

**Figure 2.**
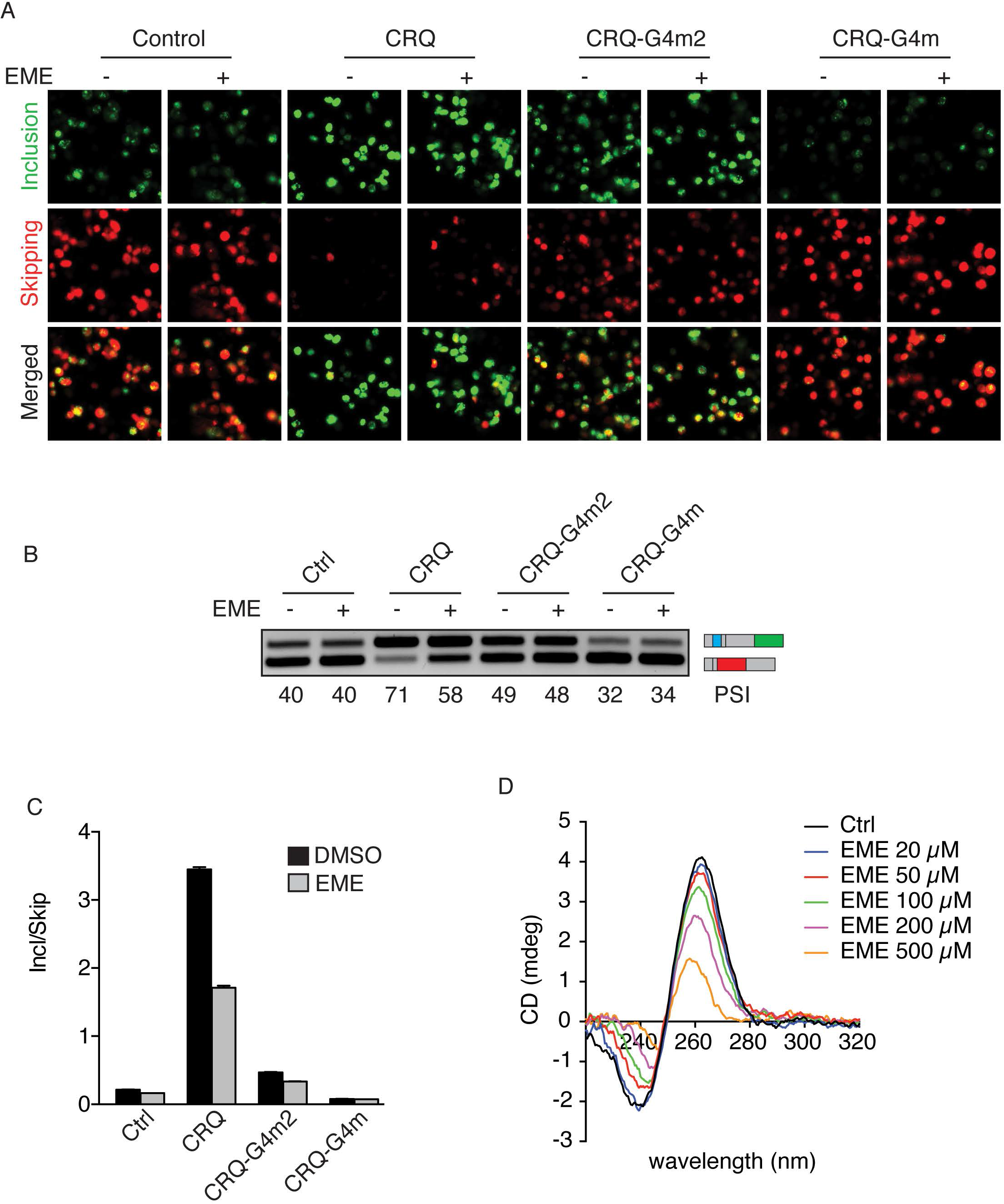
Emetine regulates alternative splicing by disrupting G-quadruplex secondary structure. (A) Fluorescence images of HEK 293FT cells transfected with CRQ and its mutants with or without the presence of emetine (20 μM). The fluorescent protein EGFP is a readout for variable exon inclusion and dsRed for variable exon skipping. (B, C) Semi-quantitative RT-PCR and qRT-PCR (C) showing that emetine promotes exon skipping only of the CRQ minigene but not control or CRQ mutant constructs whose G-quadruplex structures were destroyed. (D) CD spectra analysis of CRQ RNA oligonucleotides in the presence of different concentrations of emetine.

To further examine whether emetine directly interacts with G-quadruplex structures to disrupt quadruplex structural integrity, we conducted circular dichroism (CD) spectroscopy measurements using RNA oligonucleotides that contained the 20-nt CRQ sequences. The CD spectrum of CRQ showed a positive peak at 264 nm and a negative peak at 240 nm (Figure 2D), which is a characteristic feature of a parallel RNA G-quadruplex structure [25]. Increasing the concentrations of emetine resulted in a decrease in the CD spectra peak (Figure 2D). These results indicate that emetine treatment abolished G-quadruplexes in a dose-dependent manner.

To determine whether emetine affects additional G-quadruplexes in addition to CRQ, we examined the effect of emetine on an RNA sequence, termed I-8, that was shown to form a G-quadruplex structure (Supplementary Figure S2A) [3]. Insertion of the I-8 sequence into the minigene resulted in enhanced exon inclusion (Supplementary Figure S2B-S2D). Similar to the results observed with CRQ, emetine treatment resulted in a significant decrease in exon inclusion from 56% to 48% in the I-8 splicing minigene, but showed little effect on the I-8 mutants where the G-quadruplexes were destroyed (Supplementary Figure S2B-S2D). Moreover, CD spectroscopy measurement on I-8 RNA oligonucleotides showed that emetine perturbed the characteristic CD spectrum of the I-8 RNA in a dose-dependent manner (Supplementary Figure S2E). Together, these results further support the notion that emetine inhibits exon inclusion by disrupting G-quadruplex secondary structure.

### Cephaeline modulates alternative splicing in a G-quadruplex dependent manner

Our results suggested that the small molecule emetine inhibits G-quadruplex-dependent exon inclusion. We speculated that small molecules with a similar structure to emetine would exhibit similar properties in disrupting G-quadruplexes and regulating alternative splicing. We thus explored whether cephaeline, a closely related analog to emetine (Figure 3A), exhibits similar properties as emetine. Emetine and cephaeline differ only by the presence of one functional group, a methoxy group in emetine that is present as a hydroxyl group in cephaeline. We transfected HEK 293FT cells with the CRQ and CRQ-mutated splicing minigenes with cephaeline and observed that cephaeline treatment caused a decrease in CRQ-dependent exon inclusion from PSI 81% to 67%. By contrast, cephaeline showed minor effects on G4m2 and no effect on G4m splicing minigenes (Figure 3B-3C). In addition, cephaeline treatment also showed a decrease in the G-quadruplex containing I-8 minigene, but not the G-quadruplex mutated I-8 minigene. (Figure 3D-3E). These results indicate that cephaeline modulates alternative splicing by disrupting G-quadruplex structures in the same manner as of its analog emetine.

**Figure 3.**
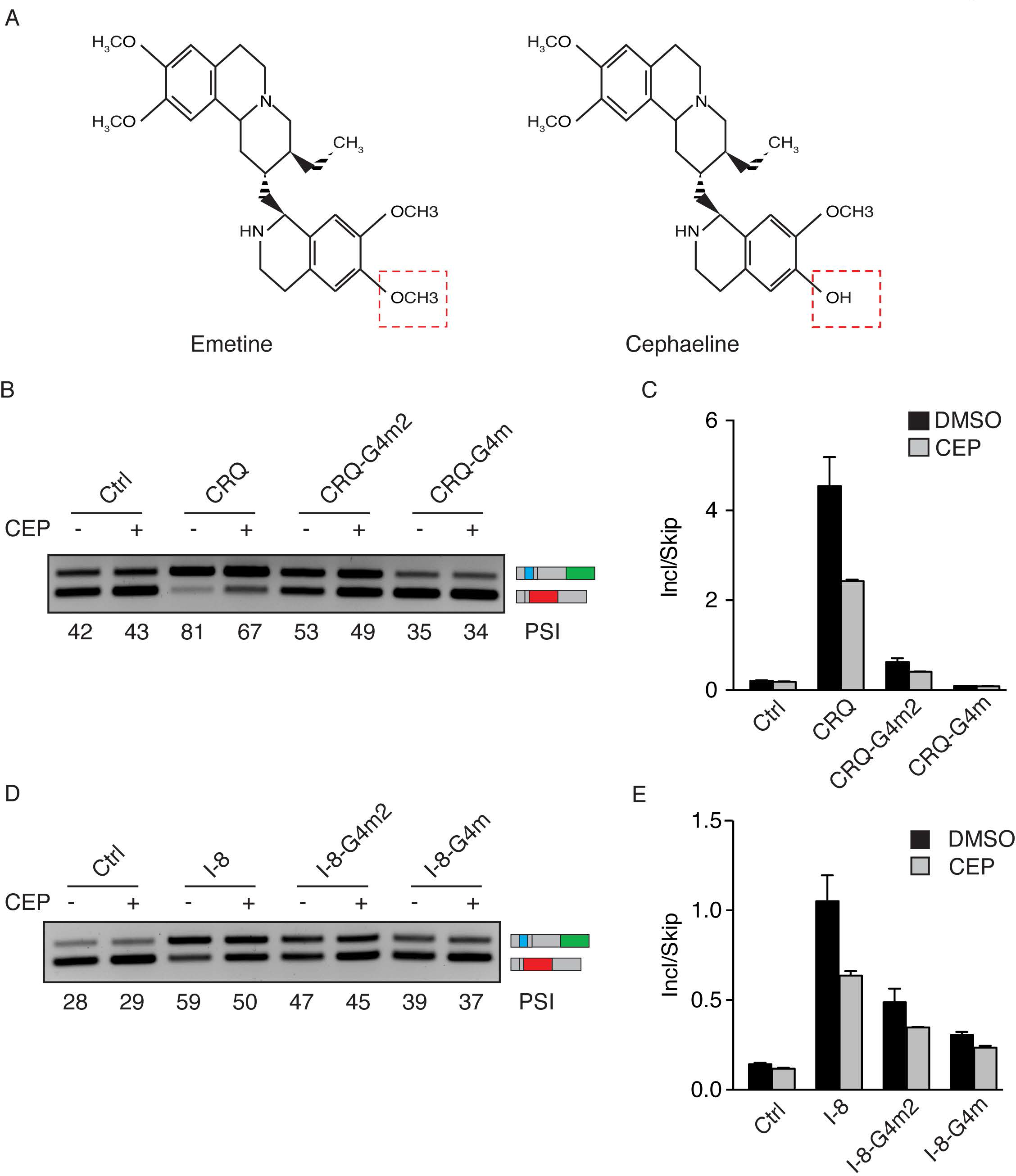
Cephaeline modulates alternative splicing on G-quadruplex-dependent manner. (A) Comparison of the chemical structures of emetine and cephaeline. The structures differ only by the moieties labeled in red boxes. (B, C) Semi-quantitative RT-PCR images (B) and qRT-PCR results (C) of HEK 293FT cells transfected with CRQ and its mutant splicing minigenes showing that cephaeline (CEP) promotes skipping of the CRQ splicing minigene and not the mutated splicing minigenes. (D, E) Effects of CEP on the I-8 G-quadruplex and its mutated splicing minigenes in HEK 293FT cells. Semi-quantitative RT-PCR images (D) and qRT-PCR results (E) are shown.

RNA G-quadruplexes are transcribed from DNA, and their DNA sequence may also form G-quadruplexes. Because transcription and splicing are coupled processes and because DNA G-quadruplexes have been shown to slow down the rate of transcription [26], we examined whether the observed RNA G-quadruplex-dependent alternative splicing was due to changes in transcription elongation. We transfected HEK 293FT cells with the splicing minigenes CRQ and its mutant CRQ-G4m and treated these cells with two transcription inhibitors, camptothecin (CPT) and 5,6-Dichlorobenzimidazole 1-β-D-ribofuranoside (DRB). Notably, CPT and DRB had little effects on the splicing of CRQ and its mutant CRQ-G4m (Supplementary Figure S3A), excluding the contribution of transcription on G-quadruplex-mediated splicing changes. Moreover, emetine has been used as a translation inhibitor [27]. To exclude the possibility that splicing changes caused by emetine were due to translation inhibition, we examined the effect of the translation inhibitor cyclohexamide (CHX) on CRQ-mediated exon inclusion. CHX exhibited little changes in alternative splicing of the CRQ minigene (Supplementary Figure S3B), illustrating that the effect of emetine on modulating alternative splicing is not due to translation inhibition.

### The effect of G-quadruplex structure on alternative splicing is not location-dependent

The above data examining the effect of disruption of G-quadruplexes on alternative splicing were conducted on constructs with the G-quadruplex located downstream of the variable exon. To test whether the location of the G-quadruplex affects alternative splicing, we introduced the same sequence of CRQ in the intron upstream of the cassette exon (CRQ-up) (Figure 4A). Transfection of the CRQ-up splicing minigene in HEK 293FT cells showed an increase in exon inclusion and decrease in exon skipping, as represented by the switch of dsRed to EGFP (Figure 4B) and by semi-quantitative RT-PCR analysis (Figure 4C, 4D). Furthermore, the CRQ-up mutants, G4m2-up and G4m-up, were also less effective in promoting exon inclusion compared to the CRQ-up construct (Figure 4B-4D). These results are consistent with the effect of CRQ inserted in the intron downstream of the cassette exon (Figure 2A-2C), indicating that the effect of G-quadruplexes on exon inclusion is not location-dependent.

**Figure 4.**
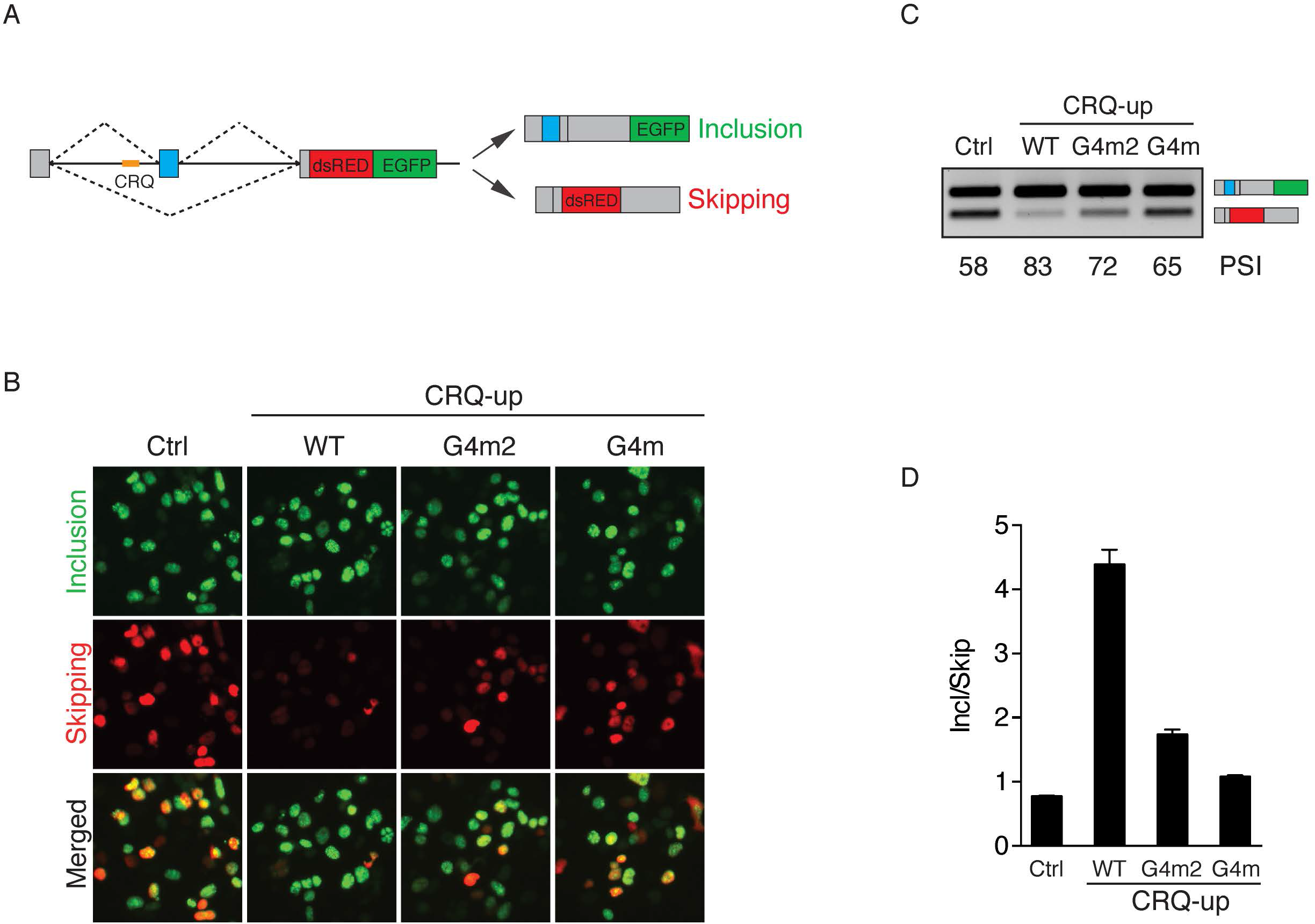
The effect of G-quadruplex structure on alternative splicing is not location-dependent. (A) Diagram of the fluorescent splicing reporter construct that contains the CRQ G-quadruplex in the intron upstream or intron downstream of the variable exon. (B) Fluorescence images of HEK 293FT cells transfected with the splicing reporters where CRQ and its mutants are inserted upstream of the variable exon. (C, D) Semi-quantitative RT-PCR images (C) and qRT-PCR results (D) of cells transfected with CRQ and its mutant minigenes showing that insertion of the CRQ G-quadruplex in the intron upstream of the variable exon stimulates exon inclusion.

We next sought to determine whether emetine inhibits exon inclusion of the CRQ-up construct. We transfected HEK 293FT cells with CRQ-up and its mutated minigenes and treated the cells with 20 μM emetine. We found that emetine promotes exon skipping only of the G-quadruplex containing CRQ-up minigene but not of the mutants (Supplementary Figure S4A-S4C), indicating that emetine specifically recognizes G-quadruplexes to alter splicing when G-quadruplexes are located in introns either upstream or downstream of the variable exon.

### G-quadruplexes are enriched near alternative exons regulated by emetine

After observing that emetine regulates alternative splicing in a G-quadruplex-dependent manner, we aimed to identify the effect of emetine on alternative splicing globally across the transcriptome. We performed RNA-sequencing using two human cell lines MCF10A and HMLE-Twist with or without emetine treatment. Emetine treatment caused significant differential splicing of thousands of splicing events in both cell lines, of which nearly 60% represented skipped exon (SE) events (Figure 5A). We focused on SE events as they were the most frequently regulated by emetine and the most common form of alternative splicing [12]. To identify a stringent set of SE regulated by emetine, we overlapped the SE events in MCF10A and HMLE-Twist and identified 1970 events shared between both cell lines and they were strongly positively correlated (pearson r = 0.89, Figure 5B). To test whether emetine mediates alternative splicing of cassette exons containing proximal G-quadruplexes, we identified all SE containing potential G-quadruplexes (PGQ) inside the cassette exon, flanking exons, or within 250 nucleotides of a splice site. We found that approximately 60 percent of emetine regulated SE contained a PGQ with 2-nt guanine tracts and approximately 10 percent contained a more stringent PGQ with 3-nt guanine tracts (Figure 5C). As examples, two of the emetine-regulated cassette exons identified as containing an exon-proximal G-quadruplex, AKR1A1 exon 7 and RFC5 exon 2, were previously identified to contain G-quadruplexes capable of regulating alternative splicing [28]. Experimental validation of these two SEs showed that emetine treatment promoted AKR1A1 exon 7 skipping and RFC5 exon 2 inclusion in both HMLE-Twist and MCF10A cells (Figure 5D, 5E). Taken together, these results illustrate that emetine regulates alternative splicing events that are enriched for G-quadruplexes globally across the transcriptome.

**Figure 5.**
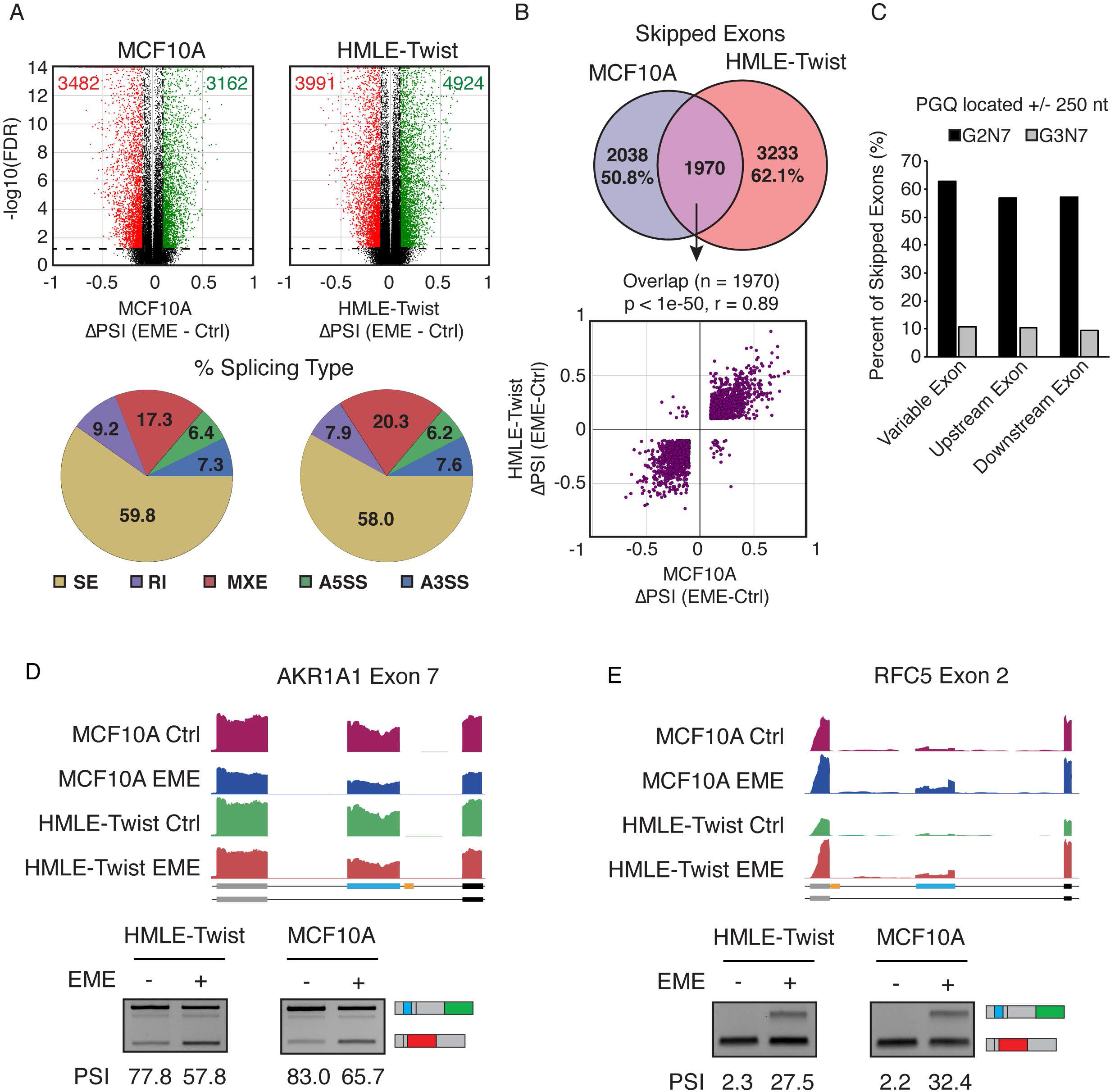
Emetine globally affects G-quadruplex associated alternative splicing. (A) Top Panel: Differential alternative splicing events identified after emetine treatment in MCF10A (Left) and HMLE-Twist (Right) cells. Alternative splicing events that show FDR ≤ 0.05 and ΔPSI ≥ 0.2 or ≤ −0.2 were colored as green and red dots, respectively. Bottom panel: Pie charts showing percentage of each splicing type represented in emetine differential splicing events. The majority of events are skipped exon (SE) events. (B) Top panel: Venn diagram showing common skipped exons regulated by emetine in both MCF10A and HMLE-Twist cells. Bottom panel: The common set of cassette exons show highly correlated regulation in both cells types. (C) Bar charts showing the percentage of skipped exon splicing events containing a G2N7 or G3N7 predicted G-quadruplex (PGQ) proximal to splice sites. (D) Genome browser tracts of RNA sequencing data and semi-quantitative RT-PCR validation showing emetine treatment promotes exon skipping of AKR1A1 Exon 7 (Left) and inclusion of RFC5 Exon 2 (Right). The G-quadruplexes are depicted in yellow in the schematics of the exon annotation.

## DISCUSSION

In recent years, emerging evidence has indicated the importance of RNA G-quadruplex structures in regulating key cellular functions. Compared to DNA G-quadruplexes which can form both antiparallel and parallel structures, RNA G-quadruplexes are less diverse in terms of strand orientation under physiological conditions and prefer to adopt a parallel-stranded structure due to the contribution of the 2’-hydroxyl group. The formation of RNA G-quadruplexes has been shown to play important roles in biological processes, such as translational regulation [29, 30], 3’ end processing [31], alternative splicing [3], and mRNA localization [32]. Despite the importance of G-quadruplexes in regulation of gene expression, few studies have investigated small molecular compounds capable of targeting these structures to denature or stabilize them.

In this study, we applied a dual-output splicing reporter containing a bona-fide G-quadruplex in a cell-based large small-molecule screen assaying for changes in alternative splicing. High-throughput screening approaches based on alternative splicing have been used in many studies [19, 33–35]. Compared to these methods, our screening system takes into account the role of RNA secondary structure on regulation of alternative splicing. In addition to the screens based on dual fluorescence signal, we applied RT-PCR validation for promising compounds, thus increasing the specificity in targeting alternative splicing and eliminating false-positive hits.

We identified emetine as a compound capable of regulating alternative splicing by interfering with the G-quadruplexes. Through mutational analysis and investigation of the G-quadruplex CD-spectra, we showed that emetine disrupts G-quadruplex secondary structure to modulate alternative splicing. These results were bolstered by similar results from our complementary analysis using cephaeline, an analog of emetine. To show that emetine functions to regulate splicing at G-quadruplexes, we examined its effects on two distinct G-quadruplex forming sequences, CRQ and I-8. Emetine has been previously identified to regulate alternative splicing of Bcl-x [36]. Interestingly, this gene contains two G-quadruplexes in its pre-mRNA [37, 38]. As a future direction, it will be interesting to investigate the contribution of these G-quadruplexes on Bcl-x alternative splicing. In addition, emetine was previously shown to inhibit translation by binding to the 40S subunit of the ribosome [24]. However, our experiments excluded the possibility that the observed splicing changes were due to translational inhibition. Although we cannot fully exclude the possibilities that emetine indirectly regulates G-quadruplex through its role as a translational inhibitor, we found that many emetine-regulated exon skipping events contain G-quadruplexes proximal to the splice sites, suggesting that emetine may have a global role in regulating G-quadruplex-dependent alternative splicing.

To date, very few studies have evaluated the interaction of small molecules with RNA G-quadruplexes. TMPYP4 has been shown to disrupt G-quadruplex structures when binding to RNA [39, 40], but it has additional effects beyond potential disruption of RNA G-quadruplexes, including poor selectivity for DNA and RNA G-quaruplexes [41] and translation inhibition [42]. Our study identifies emetine as a splicing modulatory compound. By disrupting RNA G-quadruplex structures, emetine may affect cellular functions by impacting G-quadruplex-mediated alternative splicing. Emetine can serve as a reagent to help understand the mechanisms underlying how RNA binding proteins regulate alternative splicing through the interaction with G-quadruplexes. It may also be used for the identification of splicing targets that are regulated by G-quadruplex structures. Thus, our studies may inspire researchers to consider the role of RNA secondary structures in addition to the linear sequences in regulation mechanism of alternative splicing.

Taken together, the presented study is the first to screen compounds that regulate alternative splicing by disrupting cis-elements that contain RNA G-quadruplex structures rather than a linear sequence. Our findings open new horizons for the identification of small molecules capable of regulating alternative splicing by targeting RNA secondary structures.

## ACKNOWLEDGEMENTS

We would like to thank Dr. Chi-Hao Luan and Sara Fernandez Dunne at the High Throughput Analysis Laboratory, Northwestern University for assisting the high throughput screening and analysis. We thank Dr. Jin Wang at Baylor College of Medicine for suggestions on cephaeline.

